# Investigating the genetic regulation of the expression of 63 lipid metabolism genes in the pig skeletal muscle

**DOI:** 10.1101/118950

**Authors:** Rayner González-Prendes, Raquel Quintanilla, Marcel Amills

## Abstract

Despite their potential involvement in the determination of fatness phenotypes, a comprehensive and systematic view about the genetic regulation of lipid metabolism genes is still lacking in pigs. Herewith, we have used a dataset of 104 pigs, with available genotypes for 62,163 single nucleotide polymorphisms and microarray gene expression measurements in the *gluteus medius* muscle, to investigate the genetic regulation of 63 genes with crucial roles in the uptake, transport, synthesis and catabolism of lipids. By performing an eQTL scan with the GEMMA software, we have detected 12 cis- and 18 trans-eQTL modulating the expression of 19 loci. Genes regulated by eQTL had a variety of functions such as the β-oxidation of fatty acids, lipid biosynthesis and lipolysis, fatty acid activation and desaturation, lipoprotein uptake, apolipoprotein assembly and cholesterol trafficking. These data provide a first picture about the genetic regulation of loci involved in porcine lipid metabolism.

---

The search of regulatory variants with causal effects on the expression of genes with important metabolic roles is fundamental to elucidate the genetic basis of multiple physiological and pathological phenotypes [1]. In humans, thousands of expression QTL (eQTL) have been detected so far and the majority of them appear to act locally (cis-eQTL) rather than influencing the expression of genes located at distant genomic regions or chromosomes (trans-eQTL) [1,2]. Moreover, around 50% of human cis-eQTL are shared across distinct tissues, though the consistency in the magnitude and the direction of these regulatory effects may be variable [2].

The genetic regulation of lipid metabolism genes has been poorly studied in pigs in spite of the fact that it may have a potential impact on the phenotypic variation of fatness traits. Indeed, the majority of eQTL studies performed in pigs have targeted either genes whose expression correlates with lipid phenotypes or loci comprised within the confidence intervals of fatness quantitative trait loci [3–7]. At present, we do not know if porcine lipid genes are predominantly regulated in cis- or trans- and if such regulation is featured by single or multiple polymorphisms. The goal of the current work was to shed light into these issues by identifying eQTL with effects on the muscle expression of 63 genes with an established role in the uptake, transport, synthesis and catabolism of lipids.

As animal material, we have used 104 barrows from a commercial Duroc porcine line (Lipgen population) distributed in five half-sib families. After weaning, this pig population was transferred to the experimental test station at the Centre de Control Porcí of the Institut de Recerca i Tecnologia Agroalimentàries (IRTA). A detailed description of the experimental population and management conditions has been reported [8,9]. Barrows were slaughtered at an approximate age of 190 days. *Gluteus medius* (GM) muscle biopsies were obtained in the abattoir and they were immediately frozen in liquid nitrogen, being subsequently stored at -80 °C. All animal care and management procedures followed the ARRIVE guidelines [10] and they were approved by the Ethical Committee of the Institut de Recerca i Tecnologia Agroalimentàries (IRTA).

GeneChip Porcine Genomic arrays (Affymetrix, Inc., Santa Clara, CA) were used to measure gene expression in GM samples from the 104 Duroc pigs mentioned above (data are available in the Gene Expression Omnibus public repository of the National Center for Biotechnology Information, accession number: GSE19275). Data pre-processing and normalization were carried out with the BRB-ArrayTools software version 3.7.1.[11]. Genes displaying more than 20% of expression values over ±1.5 times the median expression of all arrays were retained for further analysis. A detailed description of the techniques and methods used to perform RNA purification and microarray hybridization can be found in [12]. Finally, sixty three loci annotated in the Ensembl (S.scrofa 10.2) database and having a well established role in lipid metabolism (Supplementary Table 1) were selected for further analysis.

The Porcine SNP60K BeadChip (Illumina, San Diego, CA) was employed to genotype 62,163 single nucleotide polymorphisms (SNPs) in the 104 Duroc pigs by following a previously reported protocol [5]. The GenomeStudio software (Illumina) was employed to evaluate the quality of the typing data. By using PLINK [13], we discarded SNPs with rates of missing genotypes above 10%, minor allele frequencies (MAF) below 5%, as well as those did not conform Hardy-Weinberg expectations (threshold set at a *P*-value of 0.001). Markers that did not map to the porcine reference genome (Sscrofa10.2 assembly) and those located in sex chromosomes were also eliminated from the data set. Moreover, were eliminated SNPs that were in complete linkage disequilibrium (r^2^ > 0.98). After these filtering steps, a total of 28,571 SNPs were used to carry out a GWAS analysis for gene expression phenotypes.

**Table 1.** Cis-eQTLs regulating the expression of 10 genes involved in porcine lipid metabolism1.

| Genes |  |  | Cis-eQTL |  |  |  |  |  |  |  |  |  |
| --- | --- | --- | --- | --- | --- | --- | --- | --- | --- | --- | --- | --- |
| Symbol | SSC | Location (Mb) | SSC | N | SNP | Region (Mb) | P-value | q-value | B | $\delta \pm SE$ | A1 | MAF |
| *ACADS* | 14 | 43.1 | 14 | 34 | MARC0094155 | 42.6-45.9 | 0.00 | 0.00 | 0.00 | $-0.62 \pm 0.08$ | A | 0.21 |
| *ACOX3* | 8 | 4.3-4.4 | 8 | 1 | ALGA0118448 | 4.4 | 0.00 | 0.00 | 0.00 | $-0.75 \pm 0.18$ | A | 0.08 |
| | | | | 2 | M1GA0025674 | 2.7-3.7 | 0.00 | 0.02 | 0.03 | $-0.34 \pm 0.10$ | A | 0.36 |
| *CITED2* | 1 | 28.2 | 1 | 4 | MARC0028659 | 26.5-27.4 | 0.01 | 0.02 | 0.07 | $0.27 \pm 0.10$ | A | 0.38 |
| *HMGCS1* | 16 | 29.4 | 16 | 18 | ALGA0089927 | 28.0-29.8 | 0.01 | 0.02 | 0.14 | $0.23 \pm 0.07$ | G | 0.35 |
| *LRP6* | 5 | 63.5-63.6 | 5 | 20 | ASGA0025668 | 62.2-63.8 | 0.00 | 0.02 | 0.13 | $0.42 \pm 0.13$ | A | 0.40 |
| *LIPA* | 14 | 110.1 | 14 | 10 | ASGA0065584 | 108.8-109.9 | 0.01 | 0.04 | 0.14 | $0.27 \pm 0.11$ | G | 0.19 |
| *NCOA1* | 3 | 121.2-121.3 | 3 | 3 | MARC0003746 | 120.0-120.4 | 0.00 | 0.02 | 0.05 | $-0.28 \pm 0.08$ | A | 0.27 |
| *NPC2* | 7 | 103.5 | 7 | 3 | INRA0027651 | 104.1-104.4 | 0.00 | 0.01 | 0.04 | $0.28 \pm 0.09$ | A | 0.28 |
| | | | | 6 | ALGA0043923 | 102.5-103.1 | 0.00 | 0.00 | 0.00 | $0.33 \pm 0.07$ | A | 0.26 |
| *SLC25A17* | 5 | 4.8 | 5 | 27 | H3GA0015347 | 2.7-5.9 | 0.00 | 0.00 | 0.00 | $-0.79 \pm 0.15$ | G | 0.30 |
| *VLDLR* | 1 | 245.0 | 1 | 1 | ASGA0005756 | 244.9 | 0.00 | 0.04 | 0.04 | $-0.33 \pm 0.13$ | A | 0.20 |

<sup>1</sup>SSC: porcine chromosome, N: Number of SNPs significantly associated with traits under study, SNP: SNPs displaying the most significant associations with traits under study, Region (Mb): regions containing SNPs significantly associated with traits under study, P-value: nominal P-value, q-value: q-value calculated with a false discovery rate approach, B : Bonferroni-corrected P-value, $\delta$ : allelic effect and its standard error (SE), A1: minority allele, MAF: frequency of the minority allele.

Statistical analyses were performed with the GEMMA software [14] that uses a standard linear mixed model and an exact test of significance to identify associations between genotypes and phenotypes. The existence of population structure is taken into account by considering a relatedness matrix [14]. The model assumed in the statistical analysis was:

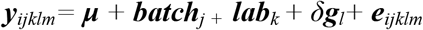

where *y_ijklm_* is the vector that describes the mRNA levels of each gene in the GM muscle of the *i^th^* individual; **μ** is the mean mRNA expression of each gene in the population; *batch_j_* and ***lab***_*k*_ are the systematic effects i.e. “batch of fattening” (with 4 categories) and “laboratory” (microarray data were produced in two distinct laboratories); *δ* is the SNP allelic effect estimated as a regression coefficient on the corresponding *g_l_* genotype (values - 1, 0, 1) of the *l^th^* SNP; and *e_ijklm_* is the residual effect. Correction for multiple testing was implemented with a false discovery rate approach [15] and SNPs with a q-value ≤ 0.05 were considered as significantly associated with gene expression. In the analysis of cis-eQTL, multiple testing was corrected by taking into consideration the number of SNPs contained within 2 Mb windows around each gene, while in the trans-eQTL analysis we took into account the whole set of 28,571 SNPs.

The eQTL scan for lipid-related genes made possible to identify 12 cis-eQTL and 18 trans-eQTL influencing the mRNA levels of 19 loci (Tables 1 and 2, Figure 1). As shown in Table 1, the two cis-eQTL detected for the *ACOX3* (SSC8: 2.7-3.7 Mb and 4.4 Mb) and *NPC2* (SSC7: 102.5-103.1 Mb and 104.1-104.4 Mb) genes were located in adjacent positions and they might correspond to two genetic determinants (instead of 4). In a previous study, Chen et al. [16] identified 120 cis-eQTLs and 523 trans-eQTLs with effects on porcine hepatic gene expression. However, they focused their study on a dataset of 300-400 genes that showed significant correlations with traits under study and their sample size was larger than ours. In the current work, the numbers of cis- and trans-eQTL for lipid genes were quite similar (Tables 1 and 2). In contrast, Cánovas et al.[12] performed a genome scan for porcine muscle expression phenotypes and observed a predominance of trans- *vs* cis-eQTL. The most likely reason for this discrepancy is that we have used different thresholds of significance to correct for multiple testing in the cis- and trans-eQTL analyses. Indeed, in humans the majority of eQTL identified so far act in cis-. For instance, a recent eQTL scan in 869 lymphoblastoid cell lines revealed that 3,534 and 48 genes were affected by eQTL in cis- and trans-, respectively [17]. Similarly, a global analysis of 53 datasets demonstrated the existence of 116,563 high confidence eQTL [18]. Around 91% and 9% of these eQTL acted in cis- and trans-, respectively [18], and there was an average of 1.8 eQTL per gene.

**Figure 1.**
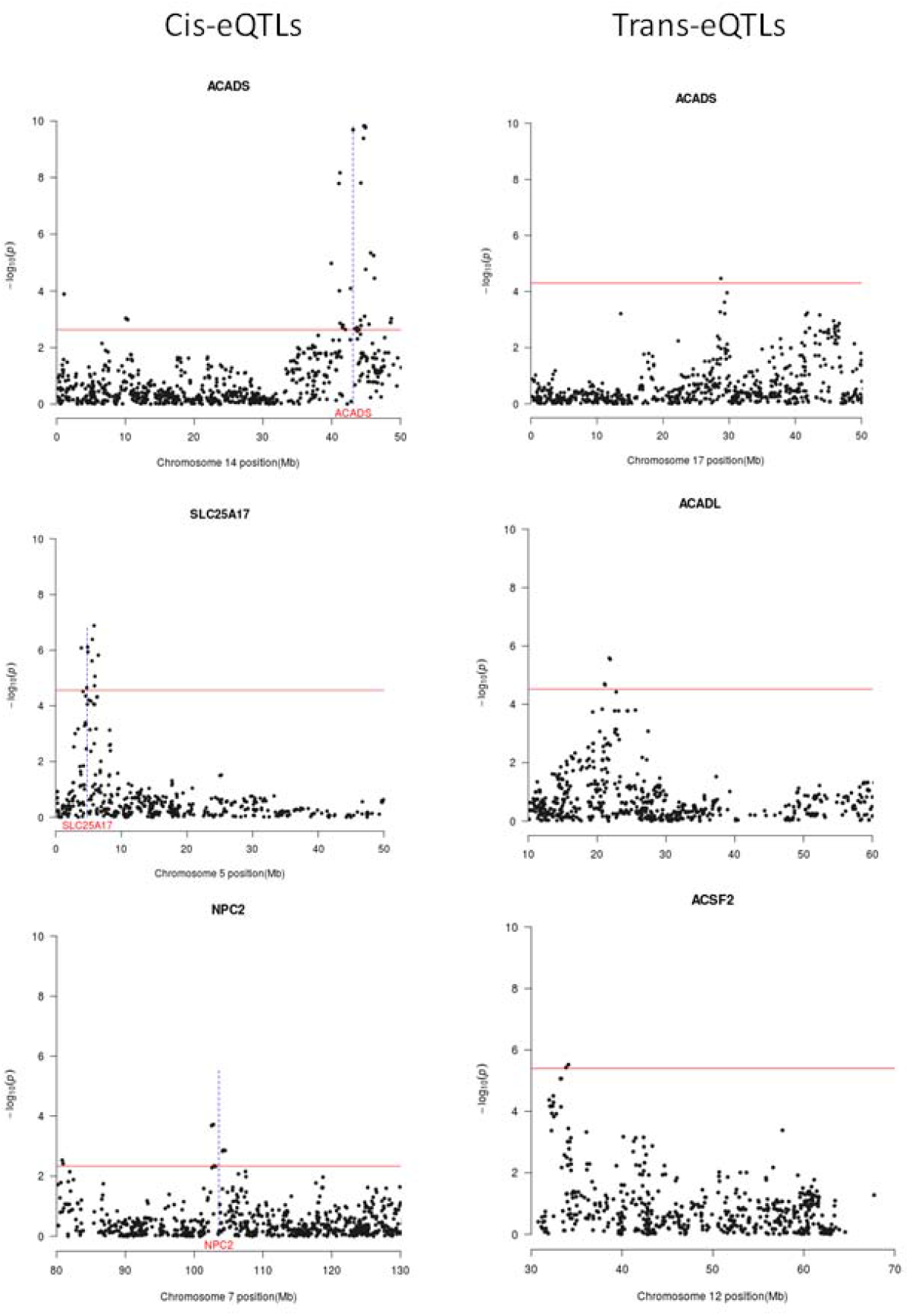
Plots of cis-eQTLs (left panel) regulating *ACADS, SLC25A17* and *NPC2* mRNA levels and of trans-eQTLs (right panel) influencing the expression of the *ACADS, ACDL* and *ACFS2* loci. The x-axis represents the chromosomal region containing the eQTL (measured in Mb), and the y-axis shows the -log10 (P-value) of the associations found. The horizontal line indicates the threshold of significance (q-value ≤ 0.05). Vertical lines in left panel plots depict the genomic location of the *ACADS, SLC25A17* and *NPC2* genes.

**Table 2.** Trans-eQTLs regulating the expression of 12 genes involved in porcine lipid metabolism^1^.

| Genes |  |  | Trans-eQTLs |  |  |  |  |  |  |  |  |  |
| --- | --- | --- | --- | --- | --- | --- | --- | --- | --- | --- | --- | --- |
| Symbol | SSC | Location (Mb) | SSC | N | SNP | Region (Mb) | P-value | q-value | B | $\delta \pm SE$ | A1 | MAF |
| *ACADL* | 15 | 124.7-124.7 | 3 | 1 | MARC0017993 | 144.3-144.3 | 0.00 | 0.03 | 0.08 | $-0.58 \pm 0.13$ | A | 0.18 |
| | | | | 2 | ALGA0123606 | 21.7-21.8 | 0.00 | 0.03 | 0.07 | $-0.58 \pm 0.13$ | A | 0.18 |
| *ACADM* | 6 | 127.5-127.5 | 13 | 7 | DIAS0003141 | 141.6-144.1 | 0.00 | 0.04 | 0.21 | $-0.72 \pm 0.15$ | A | 0.12 |
| | | | 9 | 1 | MARC0004327 | 29.5-29.5 | 0.00 | 0.04 | 0.34 | $-0.56 \pm 0.12$ | A | 0.22 |
| *ACADS* | 14 | 43.1-43.1 | 17 | 1 | INRA0053259 | 28.7-28.7 | 0.00 | 0.05 | 0.97 | $-0.46 \pm 0.09$ | A | 0.45 |
| | | | 14 | 3 | H3GA0040210 | 53.7-55.5 | 0.00 | 0.00 | 0.02 | $-0.48 \pm 0.08$ | G | 0.28 |
| | | | 12 | 3 | M1GA0017106 | 58.9-59.4 | 0.00 | 0.02 | 0.24 | $-0.54 \pm 0.12$ | G | 0.12 |
| | | | 3 | 1 | MARC0039787 | 134.6-134.6 | 0.00 | 0.01 | 0.19 | $-0.47 \pm 0.11$ | G | 0.18 |
| *ACSF2* | 12 | 26.8-26.8 | 12 | 4 | MARC0030253 | 33.2-34.0 | 0.00 | 0.01 | 0.09 | $-0.49 \pm 0.10$ | A | 0.50 |
| *APOA1* | 9 | 49.2-49.2 | 1 | 3 | MARC0004843 | 181.0-183.7 | 0.00 | 0.01 | 0.01 | $1.02 \pm 0.19$ | G | 0.07 |
| *CEBPD* | 4 | 87.3-87.3 | 7 | 1 | ALGA0045624 | 128.5-128.5 | 0.00 | 0.04 | 0.04 | $1.30 \pm 0.25$ | G | 0.04 |
| *CMIP* | 6 | 7.1-7.2 | 5 | 2 | ASGA0103424 | 12.4-12.7 | 0.00 | 0.04 | 0.36 | $0.59 \pm 0.13$ | G | 0.06 |
| | | | 13 | 7 | DIAS0003141 | 141.6-144.1 | 0.00 | 0.00 | 0.00 | $0.58 \pm 0.09$ | A | 0.12 |
| *ECI2* | 7 | 2.5-2.5 | 12 | 1 | MARC0021670 | 37.0-37.0 | 0.00 | 0.03 | 0.03 | $-0.62 \pm 0.13$ | G | 0.16 |
| *GPAT3* | 8 | 144.2-144.2 | 17 | 1 | H3GA0049617 | 61.6-61.6 | 0.00 | 0.05 | 0.18 | $0.65 \pm 0.14$ | G | 0.22 |
| *LACTB* | 1 | 120.1-120.1 | 15 | 2 | MARC0020666 | 3.2-3.4 | 0.00 | 0.05 | 0.08 | $-0.62 \pm 0.14$ | A | 0.13 |
| *LRP6* | 5 | 63.5-63.6 | 4 | 1 | MARC0056621 | 134.9-134.9 | 0.00 | 0.03 | 0.03 | $-0.42 \pm 0.08$ | G | 0.47 |
| *SLC25A17* | 5 | 4.8-4.8 | 1 | 7 | SIRI0000355 | 129.2-138.3 | 0.00 | 0.03 | 0.26 | $-0.64 \pm 0.13$ | A | 0.18 |

<sup>1</sup>SSC: porcine chromosome, N: Number of SNPs significantly associated with traits under study, SNP: SNPs displaying the most
significant associations with traits under study, Region (Mb): regions containing SNPs significantly associated with traits under study,
P-value: nominal P-value, q-value: q-value calculated with a false discovery rate approach, B : Bonferroni-corrected P-value, $\delta$ : allelic
effect and its standard error (SE), A1: minority allele, MAF: frequency of the minority allele.

The majority of trans-eQTL detected by us resided in chromosomes different than the one containing the targeted gene, suggesting that they may exert their effects through SNPs that alter the synthesis or structure of a diffusible factor. We also observed the existence of several genes *(e.g. ACADS* and *SLC25A17*) simultaneously regulated by eQTL in cis- and in trans- (Tables 1 and 2). Particularly relevant is the case of the *ACADS* gene, whose expression was modulated by one and four cis- and trans-eQTL, respectively. This finding illustrates that even simple phenotypes, such as gene expression, can be regulated in a highly complex manner.

From a functional point of view, this set of 12 cis- and 18 trans-eQTL regulated the expression of genes integrated in distinct metabolic pathways. In this way, the acyl-coenzyme A dehydrogenases for short-chain *(ACADS),* medium-chain *(ACADM)* and long-chain *(ACADL)* FA catalyse the first step in the FA β-oxidation pathway, and the enoyl-CoA delta isomerase 2 *(ECI2)* gene plays an essential role in the β-oxidation of unsaturated FA. Moreover, the solute carrier family 25 member 17 *(SLC25A17)* gene encodes a peroxisomal transporter of coenzyme-A, FAD and NAD+ cofactors [19] and it could have a role in the α-oxidation of FA [20]. We have also detected eQTL for genes comprised in lipid biosynthetic pathways (Tables 1 and 2). For instance, the glycerol-3-phosphate acyltransferase 3 *(GPAT3)* is involved in the synthesis of triacylglycerols [21], and the 3-hydroxy-3-methylglutaryl-CoA synthase 1 *(HMGCS1)* enzyme is a component of the cholesterol biosynthetic pathway [22]. Other relevant loci are the acyl-CoA synthetase family member 2 *(ACSF2)* gene, which may participate in FA activation [23], the *LACTB* gene that affects adiposity in mice females [23], the CCAAT/enhancer binding protein δ *(CEBPD)* gene that has a key role in the regulation of adipogenesis [24] and the Cbp/P300 interacting transactivator with Glu/Asp rich carboxy-terminal domain 2 *(CITED2)* locus that is involved in the regulation of hepatic gluconeogenesis[25].

## Conclusions

Our results demonstrate that around 30% of the lipid-related genes analysed in the current work are regulated by cis- and/or trans-eQTL with significant effects on their mRNA levels. In our data set, we have not detected a clear predominance of either cis- or transregulatory factors in the determination of gene expression, a result that contrasts with what has been obtained in humans where gene regulation is mostly exerted by cis-factors. In the next future, it would be worth to investigate if the set of eQTL detected herewith displays significant associations with the phenotypic variation of porcine traits of economic interest.

## Funding

Part of the research presented in this publication was funded by grants AGL2013-48742-C2-1-R and AGL2013-48742-C2-2-R awarded by the Spanish Ministry of Economy and Competitivity. We also acknowledge the support of the Spanish Ministry of Economy and Competitivity for the *Center of Excellence Severo Ochoa 2016-2019* (SEV-2015-0533) grant awarded to the Center for Research in Agricultural Genomics. Gonzalez-Prendes R. was funded by a FPU Ph.D. grant from the Spanish Ministerio de Educación (FPU12/00860). Thanks also to the CERCA Programme of the Generalitat de Catalunya. The authors are indebted to *Selección Batallé S.A.* for providing the animal material. Thanks also to Angela Cánovas for her advice in the bioinformatic analyses of gene expression.

---

**Supplementary Table 1.**
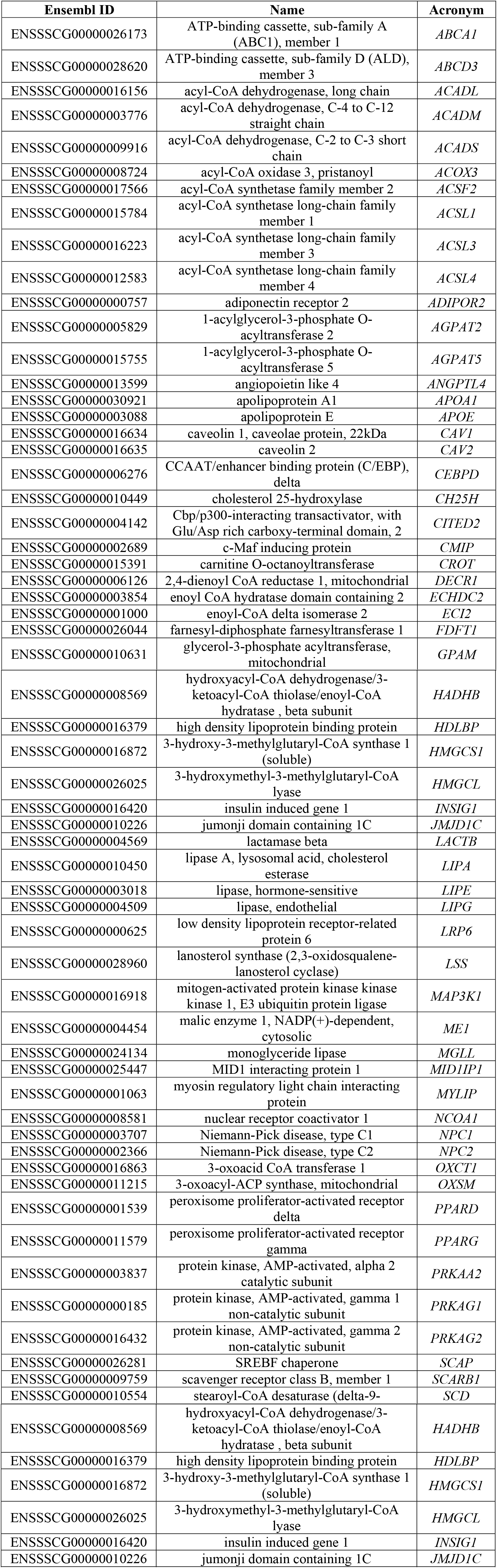
List of 63 lipid-related genes analysed in the current work.

